# A novel nuclear genetic code alteration in yeasts and the evolution of codon reassignment in eukaryotes

**DOI:** 10.1101/037226

**Authors:** Stefanie Mühlhausen, Peggy Findeisen, Uwe Plessmann, Henning Urlaub, Martin Kollmar

## Abstract

The genetic code is the universal cellular translation table to convert nucleotide into amino acid sequences. Changes to sense codons are expected to be highly detrimental. However, reassignments of single or multiple codons in mitochondria and nuclear genomes demonstrated that the code can evolve. Still, alterations of nuclear genetic codes are extremely rare leaving hypotheses to explain these variations, such as the ‘codon capture’, the ‘genome streamlining’ and the ‘ambiguous intermediate’ theory, in strong debate. Here, we report on a novel sense codon reassignment in *Pachysolen tannophilus*, a yeast related to the Pichiaceae. By generating proteomics data and using tRNA sequence comparisons we show that in *Pachysolen* CUG codons are translated as alanine and not as the universal leucine. The polyphyly of the CUG-decoding tRNAs in yeasts is best explained by a *tRNA loss driven codon reassignment* mechanism. Loss of the CUG-tRNA in the ancient yeast is followed by gradual decrease of respective codons and subsequent codon capture by tRNAs whose anticodon is outside the aminoacyl-tRNA synthetase recognition region. Our hypothesis applies to all nuclear genetic code alterations and provides several testable predictions. We anticipate more codon reassignments to be uncovered in existing and upcoming genome projects.

## Introduction

The genetic code determines the translation of nucleotide into amino-acid sequences. Any change in the code altering the meaning of a codon would introduce errors into every translated message and thus be highly detrimental or lethal (Knight et al. 2001a). Therefore, negligible if being optimal or not (Freeland and Hurst 1998; Freeland et al. 2000) the canonical genetic code was long thought to be immutable and termed a "frozen accident" of history (Crick 1968). However, reassignments of single or multiple codons in mitochondria (Knight et al. 2001b) and nuclear genomes (Lozupone et al. 2001; Miranda et al. 2006) demonstrated that the code can evolve (Knight et al. 2001a; Koonin and Novozhilov 2009; Moura et al. 2010). Several codons have been reassigned in independent lineages. Most nuclear code alterations reported so far are stop codon and CUG codon reassignments. Mainly three theories have been proposed to explain reassignments in the genetic code. The *codon capture* hypothesis states that a codon and subsequently its then meaningless cognate tRNA must disappear from the coding genome, before a tRNA with a mutated anticodon appears changing the meaning of the codon (Osawa and Jukes 1989; Osawa et al. 1992). Genome replication bias (GC or AT pressure) is thought to cause codon disappearance. In contrast, the *ambiguous intermediate* hypothesis postulates that either mutant tRNAs, which are charged by more than one aminoacyl-tRNA synthetase, or misreading tRNAs drive genetic code changes (Schultz and Yarus 1994, 1996). The ambiguous codon decoding leads to a gradual codon identity change that is completed upon loss of the wild-type cognate tRNA. The alternative CUG encoding as serine instead of leucine in *Candida* and *Debaryomyces* species (the so-called alternative yeast code; AYCU) has strongly been promoted as example for the *ambiguous intermediate* theory. Stated reasons are that the CUG codon decoding is ambiguous in many extant *Candida* species (Tuite and Santos 1996; Suzuki et al. 1997), the CUG codon decoding can-at least in part-be converted (Santos et al. 1996; Bezerra et al. 2013), and the origin of the 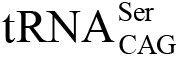 has been estimated to precede the separation of the *Candida* and *Saccharomyces* genera by about 100 million years (Massey et al. 2003). However, the ambiguous decoding of the CUG triplet in extant "CTG clade" species is caused by slightly inaccurate charging of the 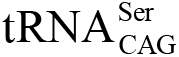 and not by competing tRNAs. Third, the *genome streamlining* hypothesis notes that codon changes are driven by selection to minimize the translation machinery (Andersson and Kurland 1995). This best explains the many codon reassignments and losses in mitochondria. In Saccharomycetaceae mitochondria, for example, ten to twenty-five sense codons are unused and the CUG codons are usually translated as threonine through a tRNA^Thr^, which has evolved from a tRNA^His^ ancestor (Su et al. 2011). In the Eremothecium subbranch, the CUG codons are decoded by alanine, but this modified code did not originate by capture of the CUG codons through an anticodon-mutated tRNA^Ala^ but by switching the acceptor stem identity determinants of the Saccharomycetaceae tRNA^Thr^ from threonine to alanine (Ling et al. 2014).

By analysis of the conservation of amino acid types and CUG codon positions in motor and cytoskeletal proteins we recently showed that these proteins allow to unambiguously assign the standard genetic code or AYCU to yeasts (Mühlhausen and Kollmar 2014). Plotting the assigned code onto the yeast phylogeny demonstrated the AYCU to be polyphyletic with "CTG clade" species and *Pachysolen tannophilus* grouping in different branches. Sequence conservation clearly showed that *Pachysolen* does not encode CUG by leucine. However, *Pachysolen* did not share any CUG positions with other yeasts. This prompted us to determine the identity of the *Pachysolen* CUG encoding by molecular phylogenetic and proteome analyses.

## Results

### A new nuclear genetic code in the yeast *Pachysolen tannophilus*

We determined the tRNA_CAG_s in 60 sequenced yeast species (Table S1) and aligned them against known 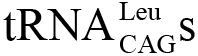 and 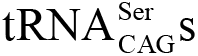. While 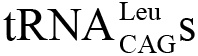 and 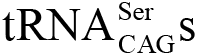 could clearly be classified, the identified *Pachysolen* tRNA_CAG_ sequence was dissimilar to both (Fig. S1). Comparison to all *Candida albicans* cytoplasmic tRNAs suggested close relationship to alanine tRNAs. The alanine identity of the *Pachysolen* tRNA_CAG_ was verified by molecular phylogenetic analyses based on extensive sequence and taxonomic sampling (Fig. 1A, Figs. S1-S5, Table S1). The sequence identity of the *Pachysolen* tRNA_CAG_ to all identified yeast GCN-decoding tRNAs (752 Saccharomycetales clade sequences) is on average 69.3%, ranging from 62.1 to 77.0%. This is slightly below the average 83.6% sequence identity within the GCN-decoding tRNAs (minimum sequence identity is 50.6%) reflecting some sequence divergence beyond the anticodon (Fig. S2). The major general alanine tRNA identity determinants, the discriminator base "A73” and the invariant "G3:U70” wobble base pair as part of the conserved 5’ sequence G^1^GC^4^ (Musier-Forsyth et al. 1991; Saks et al. 1994; Giegé étal. 1998; Giegé and Eriani 2015), are also present in the *Pachysolen* tRNA_CAG_ (Fig. S1). In contrast, serine tRNAs are characterized by a conserved variable loop sequence, and leucine tRNA_nagS_ by invariant A35 and G37 nucleotides (Fig. S1) (Saks et al. 1994; Giegé étal. 1998; Giegé and Eriani 2015). However, in *E.coli* not the anticodon sequences but different tertiary structures seem to be important for discriminating serine and leucine tRNAs (Asahara et al. 1993,1994). Although identity elements are invariant for the respective codon family box tRNAs, the same elements might be present in other tRNAs where they are located in variable regions. For example, the "G3:U70” wobble base pair, which is usually only found in alanine tRNAs, is also present in Phaffomycetaceae and some Saccharomycetaceae 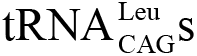, although these tRNAs clearly belong to the CTN codon box family tRNAs (Fig. 1A and Figs. S2 to S5) and CUG codons are translated as leucine (Mühlhausen and Kollmar 2014). The presence of G37 in most *Candida* 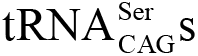 also causes partial mischarging by leucine-tRNA synthetases (Suzuki et al. 1997). The anticodon loop of the *Pachysolen* tRNA_CAG_, including the conserved A3 5 and G37 nucleotides, is identical to leucine tRNAs, while the rest of the sequence is similar to alanine tRNAs (Fig. SI).

**Figure 1:**
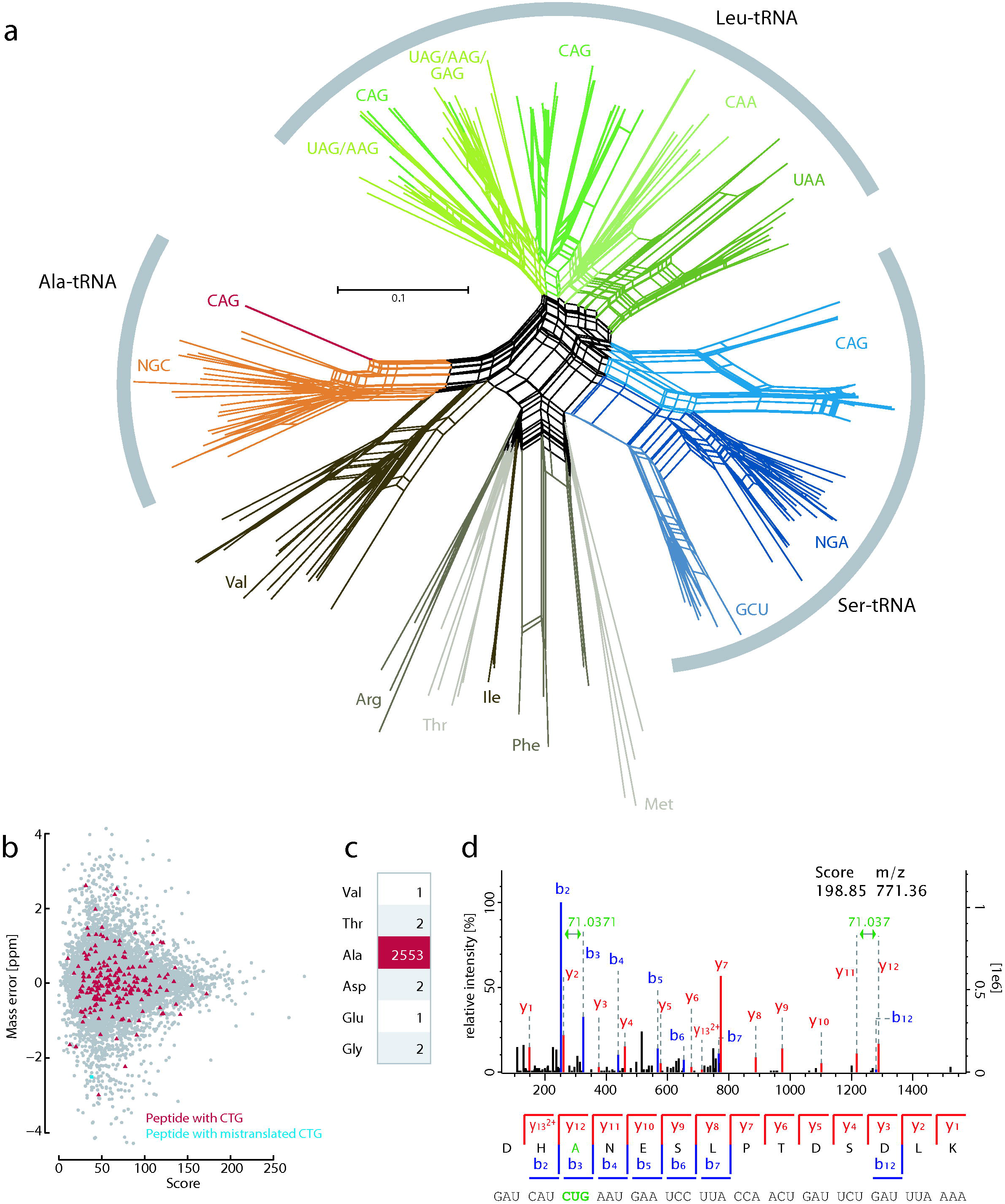
*Pachysolen* translates CUG codons with alanine instead of the universal leucine. a) Unrooted phylogenetic network of 172 tRNA sequences as generated using the Neighbour-Net method as implemented in SplitsTree v4.1.3.1. Methionine, isoleucine, arginine, and threonine tRNAs were included as outgroup. Leu-, Ser-, and Ala-tRNA_CAG_s are highlighted in dark green, blue, and red, respectively. b) Distribution of the mass errors of randomly selected 10% of all peptides (grey dots), peptides with CUG translated as alanine (red), and peptides with CUG translated as other amino acids (blue). c) Number and distribution of observed CUG translations in all peptide spectrum matches from MS/MS analysis. d) Representative MS/MS spectrum of a peptide containing CUG translated as alanine. The peptide sequence is shown below the spectrum, with the annotation of the identified matched N-terminal fragment ions (b-type ions) in blue and the C-terminal fragment ions (y-type ions) in red. Only major identified peaks are labeled for clarity (full annotation is shown in Fig. S7).

### Unambiguous translation of the CUG codons by alanine

To verify translation of the CUG codons by alanine we analysed a cytoplasmic extract of laboratory grown Pachysolen by high-resolution mass spectrometry (LC-MS/MS) generating approximately 460,000 high-quality tandem mass spectra. Spectra processing resulted in 27,126 non-redundant peptide matches with a median mass measurement error of about 240 parts per billion (Fig. 1B and Fig. S6). We identified 54% (2,844) of the 5,288 predicted proteins with median protein sequence coverage of approximately 20%, and median numbers of peptides and corresponding PSMs identified per protein of 6 and 9, respectively (Fig. S6). Of the non-redundant peptides, 666 contained sequences with CUG codons fully supported by b-and/or y-type fragment ions. Almost all of these (99.7%) contain the CUG codons translated as alanine (Figs. 1C and 1D, Fig. S7). None of the identified CUG codons are translated as serine or leucine. The small number of CUG codons translated into other amino acids might be due to differences between our and the sequenced *Pachysolen* strain (Liu et al. 2012), transcription and translation errors, or might represent spurious mischarging of the *Pachysolen* 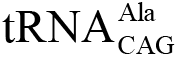. Substantial mischarging of *Candida* 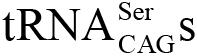 by leucines has been shown *in vitro* and *in vivo* (Suzuki et al. 1997), but other potential mischargings have never been analysed. We suppose that genetic code changes such as the CUG translation as alanine in *Pachysolen* also remained undetected in the analyses of other sequenced genomes.

### History of the CUG-decoding tRNA

To reconstruct the history and origin of all yeast tRNA_CAG_s we performed in-depth phylogenetic analyses of all UCN-decoding tRNAs (serine), GCN-decoding tRNAs (alanine), and CUN-decoding tRNAs (leucine; Figs. S8-S10). These analyses support our previous assumption (Mühlhausen and Kollmar 2014) that all "CTG clade" species' tRNA_CAG_s are serine-tRNAs, and that all Saccharomycetaceae, Phaffomycetaceae, Pichiaceae, and Kuraishia species' tRNA_CAG_s are leucine-tRNAs. Monophyly of the 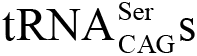 indicates a common origin in the ancestor of the "CTG clade". The UCN-decoding tRNAs split into two major subbranches, a group of UCG-decoding tRNAs to which the 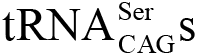 belong, and a group of UCU-decoding tRNAs. This supports a previous notion (Massey et al. 2003) that the 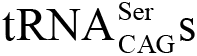 rather originated from an UCG-tRNA by insertion of an A into the anticodon than from an UCU-tRNA by insertion of a C directly before the anticodon. Saccharomycetaceae UCA-decoding tRNAs were derived from a UCG-decoding tRNA by duplication and change of anticodon, and Phaffomycetaceae, "CTG clade" species, Pichiaceae, and Kuraishia species' UCA decoding tRNAs have common ancestors with UCU-decoding tRNAs. In contrast to the monophyletic 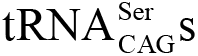, the 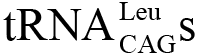 are polyphyletic and many yeasts contain several 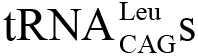 derived from gene duplication of cognate and isoacceptor tRNAs (Fig. 2A). For example, *Yarrowia lipolytica* contains thirteen 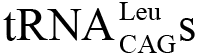, two most probably derived from either an ancestral 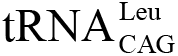 or 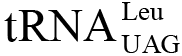 and eleven derived by gene duplication and mutation from a 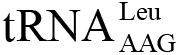 The Phaffomycetaceae, Kluyveromyces, *Lachancea kluyveri* and *Eremothecium* 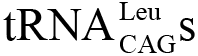 were derived from an ancestral 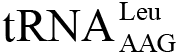, the *Lachancea thermotolerans* and *Lachancea waltii* 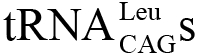 were derived by a recent gene duplication and anticodon mutation from 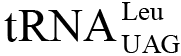, and the Pichiaceae and Kuraishia species all have derived their 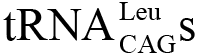 by species-specific events (Fig. 2). The CUN family box tRNAs are split into two major groups, a group of 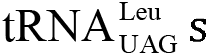 and a group of 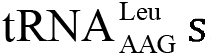. The tRNA_gagS_ are only present in Saccharomycetacea and have been derived from a 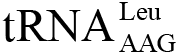. The 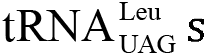 form two subgroups, which most probably originated after the split of *Yarrowia lipolytica.* One of the subgroups is restricted to *Yarrowia, Pachysolen* and the Pichiacea branches, the other is common to all yeasts and contains the *Saccharomyces* 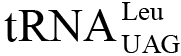. The latter is unique because it is the only *Saccharomyces cerevisiae* tRNA with an unmodified uridine in the wobble position of the anticodon triplett (Randerath et al. 1979; Johansson and Byström 2005). Unmodified U34s have otherwise only been found in mitochondria, chloroplasts and *Mycoplasma* species. The *Saccharomyces* 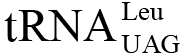 is able to translate all six leucine codons (Weissenbach et al. 1977). In contrast, modifications to U34, such as 5’-methoxycarbonylmethyl-2-thiouridine in Gln-, Lys-, and Glu-decoding tRNAs in most if not all prokaryotic and eukaryotic species, often restrict codon recognition to codons ending in A (Johansson et al. 2008; Rezgui et al. 2013). As a consequence, all sequenced yeasts have distinct NNA-decoding and NNG-decoding tRNAs for all respective two-codon families and most four-codon families. However, while all Saccharomycetaceae and Phaffomycetaceae contain this unique 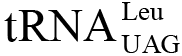, this subbranch tRNA is mutated to 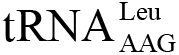 or 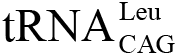 in *Pachysolen* and all Pichiaceae (Fig. 2A). The reasons for the unmodified U34 in the *Saccharomyces* 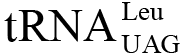 are unknown, but it is tempting to assume that the ancient yeast 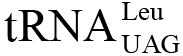 contained a modified U34 preventing ambiguous decoding of the CUG codons. If the *Yarrowia-, Pachysolen-* and Pichiacea-specific 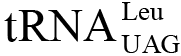 contained a modified U34, the decoding of the CUG codons in the extant species of these branches would not be ambiguous, which is supported by the unambiguous decoding of the CUG codons by alanine in *Pachysolen.*

**Figure 2:**
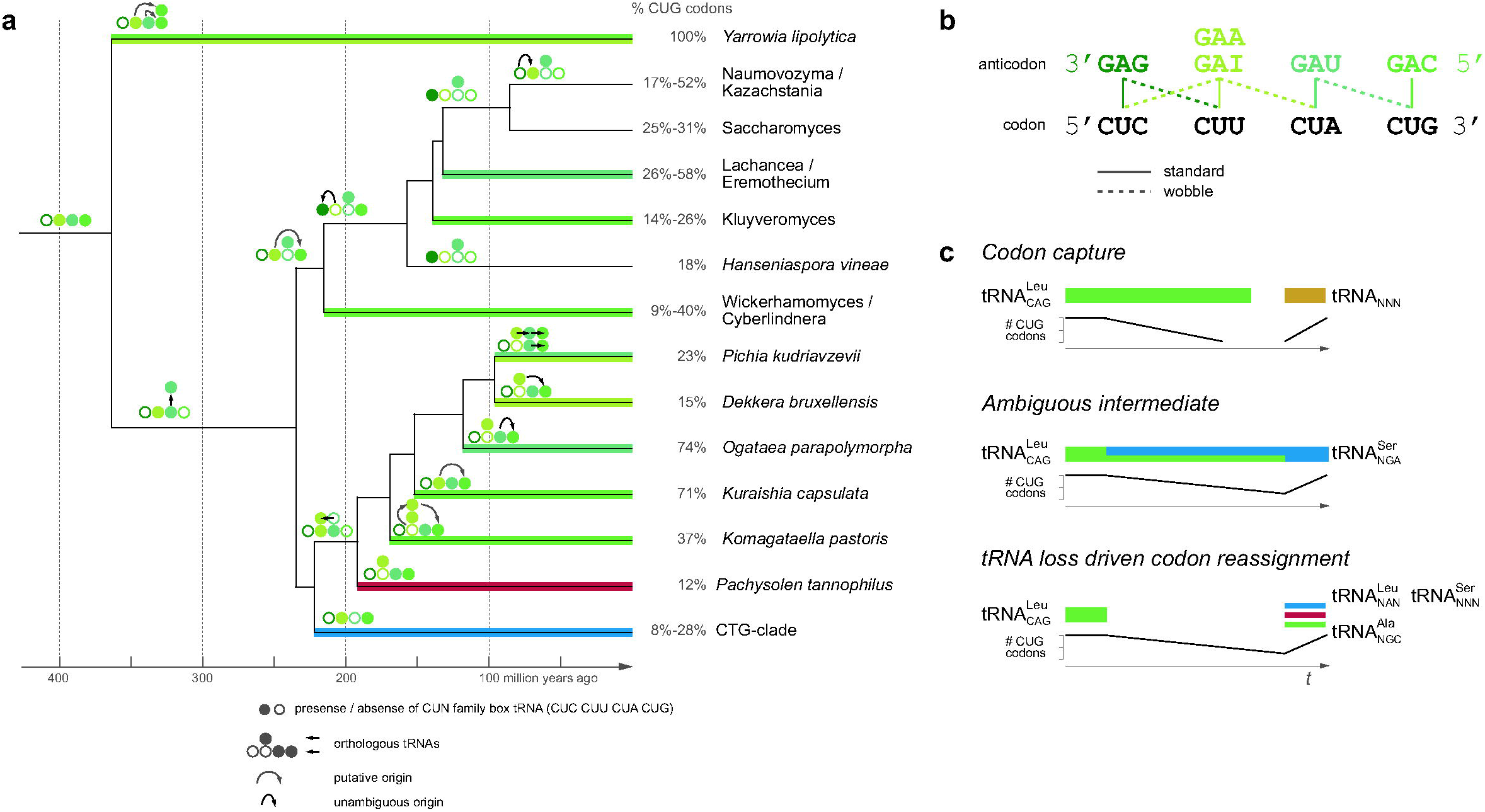
The mechanism of CUG codon reassignment. a) Decoding and origin of the tRNA_CAG_s were plotted onto a time-calibrated yeast phylogeny adapted from (Mühlhausen and Kollmar 2014) using the same colour scheme as in Fig. 1A. Coloured lines denote the presence of a tRNA_CAG_ with the colour representing the origin of the respective tRNA_CAG_. For example, *Yarrowia* contains thirteen 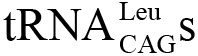, of which two most probably derived from either an ancestral 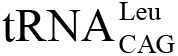 or 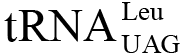 (mantis green line) and eleven most probably derived from an ancestral 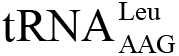 (yellow green line), and *Pachysolen* contains a 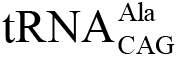 (red line). Branches with similar decoding schemes have been collapsed. Filled and empty circles at branches denote the presence and absence, respectively, of the CUN family box tRNAs, and arrows indicate tRNA gene duplications followed by anticodon mutations. The percentages of CUG codons per species and taxon were derived from (Mühlhausen and Kollmar 2014). b) The four CUN codons can be decoded by several combinations of tRNAs using standard and wobble base pairing. c) The scheme contrasts the presence of tRNA_CAG_s and number of CUG codons according to the *tRNA loss driven codon reassignment* hypothesis with assumptions based on the *codon capture* and *ambiguous intermediate* theories.

## Discussion

### Assessing models for genetic code evolution

How did such diversity in origin and decoding (leucine versus serine versus alanine) evolve? Already many years ago, several testable predictions of each of the codon reassignment hypotheses have been summarized (Knight et al. 2001a). These, however, did not include the appearance of a yeast Ala-tRNA_CAG_. Does the presence of the 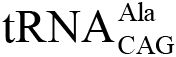 still fit into existing models? According to the *codon capture* theory, CUG codons need to have disappeared before their reassignment at the split of the "CTG clade" (Fig. 2). The time frame for codon disappearance is defined by the splits of Saccharomycetaceae / Phaffomycetaceae and *Pachysolen* / "CTG-clade" branches (hereafter called *SP* and *PC* branches, respectively). Codon disappearance must have happened either in the very short time frame between the split of the *SP* and *PC* branches and the divergence of the "CTG clade”, or before the *SP*-*PC* split, which would then necessarily include re-appearance of the 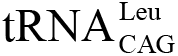 and CUG codons in the *SP* branch. Subsequently, the Ala-and Leu-tRNA_CAG_s could have captured the still unassigned CUG codon in the *Pachysolen* and the Pichiacea branches independently from each other. However, disappearance of an entire codon from a genome by neutral mutations is extremely unlikely to happen in short time and it is unlikely that genome replication bias caused only one codon to disappear. The *ambiguous intermediate* theory assumes the presence of both the cognate 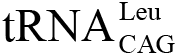 and the new 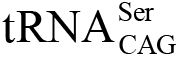 before the split of the *SP* and *PC* branches (Massey et al. 2003). It is highly unlikely that the 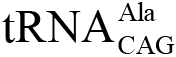 had been present at the same time giving rise to an even more ambiguous decoding by three tRNAs. More likely seems a scenario including two successive ambiguous intermediates. In this scenario, a time span with another ambiguous CUG codon decoding (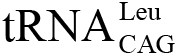 competing with 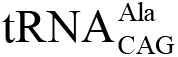) would have followed the split of the "CTG clade". If the *ambiguous intermediate* theory were true, 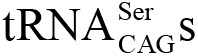 and 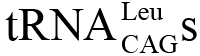 in extant species should have been derived from the same ancestral Ser-and Leu-tRNAs. While this is true for the 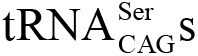 of the "CTG clade", the 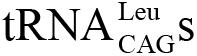 are polyphyletic and clearly have different origins (Fig. 2, Figs. S8 and S9). In addition, the *ambiguous intermediate* theory would at least require-in case of the more likely separate ambiguous intermediate events-the independent loss of the 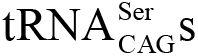 in the *SP* and Pichiacea / *Pachysolen* branches *{PP* branch), the loss of the 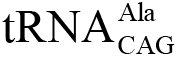 in the Pichiacea branch as part of the second ambiguous intermediate event, and the independent loss of the 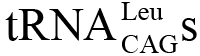 in multiple branches including the branches with altered decoding. The *ambiguous intermediate* theory does also not explain that the same codon became ambiguous in two subbranches of the same taxon in very short time.

### The *tRNA loss driven codon reassignment* mechanism

The observed polyphyly of the CAG-tRNAs and the CTN family box reassignments are better described by a *tRNA loss driven codon reassignment* process as follows (Fig. 2): The Saccharomyces type 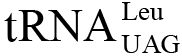 containing the unmodified U34 appeared by gene duplication of a "normal” 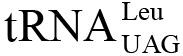 before or after divergence of *Yarrowia* enabling decoding of the CUG codons by wobble base pairing (Crick 1966). As a consequence or as an independent event the ancestor of the *SP* and *PC* clades lost its 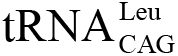 by gene loss or mutation. This tRNA loss might not have caused any viability issues because CUG codons could still be decoded as leucine by wobble base pairing (Crick 1966) by the U34-unmodified 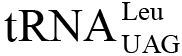 or by, although rather inefficiently, wobble base pairing by the U34-modified 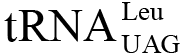. Subsequently, reduction in translational fidelity (Gromadski et al. 2006) might have caused the number of CUG codons to gradually decrease. Compared to the most ancient yeast species *Yarrowia*, all analysed yeasts have considerably decreased numbers of CUG codons (Fig. 2) (Mühlhausen and Kollmar 2014). Even the highly GC-rich genomes of *Ogataea parapolymorpha* (Ravin et al. 2013) and *Kuraishia capsulata* (Morales et al. 2013) have less CUG codons suggesting a general strong reduction of CUG codon usage after the divergence of *Yarrowia.* Increased AT pressure has been suggested as reason for reduced GC content at the N3 codon position in "CTG clade” species (Massey et al. 2003), and a similar AT bias is also present in coding genes of *Pachysolen.* However, AT pressure would have applied on all codons and cannot explain the observed reduction in CUG codon usage in all yeasts alone. CUG codon reduction most probably happened by transitions to other leucine codons. Within protein coding regions codon changes within the CTN family box, and also between CTG and TTG, are extremely frequent, demonstrated by the observation that not even closely related *Saccharomyces* species have considerable numbers of conserved leucine codons (Mühlhausen and Kollmar 2014). The unassigned CUG codon could have subsequently been captured by other tRNAs with mutated anticodons. Thus, in the "CTG clade" a mutated Ser-tRNA could capture the CUG codon triggered or supported by the loss of the 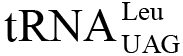, which eliminated the possibility of further ambiguous CUG decoding. In the *PP* branch, *Pachysolen* subsequently acquired the 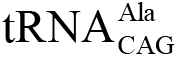 from one of the GCN-decoding tRNAs, and the Pichiaceae and Kuraishia species captured 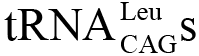 independently from each other (Figs. 2A and S8). In principle, it is highly unlikely that sense codons can be captured by tRNAs charged with noncognate amino acids because of the high recognition accuracy of the respective tRNAs by the aminoacyl-tRNA synthetases (Crick 1968; Saks et al. 1994). Aminoacyl-tRNA synthetases (aaRSs) recognize their cognate tRNAs at least at two different regions, most prominently at the discriminator nucleotide "N73" at the acceptor stem, and at the anticodon (Saks et al. 1994; Giegé et al. 1998; Giegé and Eriani 2015). However, the recognition sites of leucine, serine and alanine aaRSs do not include the respective anticodons of leucine, serine, and alanine codons, providing an explanation that only these tRNAs (in addition to any leucine tRNA) could capture the free CUG codon. Changing the anticodons of other tRNAs would disrupts charging by the respective aaRSs and thus prevents capture of noncognate codons. The *codon capture* theory distinguishes itself from the *tRNA loss driven codon reassignment* mainly by the order of events (tRNA loss after or before the reduction of CUG codons, respectively), the degree of CUG codon loss (loss of all codons versus a reduction of CUG codon usage, respectively) and the cause of reduction of CUG codons (AT pressure versus tRNA loss and decreased decoding fidelity, respectively; Fig. 2C). In contrast to the *ambiguous intermediate* hypothesis, the *tRNA loss driven codon reassignment* does not require any ambiguous decoding of the CUG codons. In addition, capture of the CUG codon by different tRNAs is an elemental characteristic of the *tRNA loss driven codon reassignment* and does not need to be split into independent events.

### Extending the new hypothesis to the evolution of other codon reassignments

Similar to CUG codon reassignment, our hypothesis can be applied to the reassignment of codons in other nuclear codes such as the stop codons in ciliates and diplomonads (Knight et al. 2001a; Lozupone et al. 2001). Reassignment in these cases might have happened by mutation of the single eukaryotic release factor eRF1 freeing respective stop codons. In contrast to bacteria that often have polycistronic mRNAs, eukaryotic mRNAs are usually monocistronic. Thus, read through of eukaryotic mRNAs does not impose any consequences other than elongation of protein tails by few residues. Examples are the UAA and UAG codons that were captured by single-base mutated Gln-and Glu-tRNAs. Both cognate tRNAs and the tRNAs with stop codons can still be charged since the glutamyl-tRNA synthetase does not discriminate the third position of the anticodon (Nureki et al. 2010). Another example supporting the *tRNA loss driven codon reassignment* hypothesis is *Mycoplasma capricolum*, which lacks a dedicated tRNA for decoding CGG but still contains six CGG codons in its genome (Oba et al. 1991). In contrast to the expectations of the *codon capture* theory, the tRNA is lost but codons are still present, which could, although inefficiently, be translated through wobble base pairing by the CGA-decoding tRNA. For genomes with unused codons such as *Micrococcus luteus*, which lacks AUA and AGA codons (Kano et al. 1993), and also mitrochondria, of which most if not all lack some codons, it is often not obvious whether tRNA loss or codon loss happened first. In genomes, which lack entire codon boxes, codon loss is more likely to have preceded tRNA loss, while tRNA loss most likely happened before codon disappearance of single codons of codon box families.

In mitochondria, codon reassignments have happened rather frequently. However, mitochondrial genomes are extremely small and changes in decoding only affect a few genes. In contrast to most nuclear genomes, where four-codon families require at least two tRNAs, most mitochondria require only one tRNA with a U in the wobble position for all four codons. Therefore, reassignments mainly affect codons in mixed codon boxes such as stop codons, AUA isoleucine, AGR arginine, and AAA lysine. These reassignments might have happened according to the *tRNA loss driven codon reassignment* hypothesis because all mentioned codons could be translated, although very inefficiently, by another tRNA of the codon family box. Usually, this ambiguous decoding is not competitive and happens only occasionally. However, it preserves the organism’s viability after loss of one of the respective tRNAs. For example, the TGA stop codon in mitochondrial genomes can be translated by the tryptophane íRNACCA, which has also been shown to happen at low levels under selective pressure in *Bacillus subtilis* (Lovett et al. 1991). Subsequent to tRNA loss, the inefficient capturing tRNAs are often optimized for decoding both the cognate and the captured codon by mutation (e.g. the tryptophane tRNAs in many mitochondria, where they also decode TGA codons, have a C to U mutation in the wobble position of the anticodon creating a UCU that can pair with both UGA and UGG) and/or by nucleotide modification such as 5’-taurinomethylation of the wobble uridine in mitochondrial methionine íRNAAUR (Suzuki et al. 2011; Watanabe and Yokobori 2011) and lysidine and agmatidine modification of the wobble C in isoleucine íRNACAU in bacteria and archaea, respectively (Voorhees et al. 2013). Instead of adjusting the tRNA for higher decoding fidelity, the aaRS might become adapted by compensatory mutations to slightly different amino acid identity patterns. This could happen by enabling two different tRNAs be charged by the same aaRS, or by gene duplication and subsequent adaptation of the two aaRSs to the respective tRNAs. An example for the latter case is the decoding of the CUN codons in yeast mitochondria by threonine. The threonine íRNAUAG originated from a histidine tRNAcuc (Su étal. 2011) and requires a dedicated ThrRS for proper charging (Pape et al. 1985). In *Eremothecium*, this íRNAUAG was mutated to "G3:U70” (Ling et al. 2014) and is accordingly charged by the AlaRS, which is non-discriminative against the anticodon. All these examples support our assumption that only tRNAs charged by aaRS that are non-discriminative against the mutated anticodon bases can subsequently capture free codons. However, there are a few examples for reassignments where a change of a discriminator base should have disfavoured correct charging. For example, UGA is translated as glycine in SR1 bacteria from the human microbiota (Campbell et al. 2013), although both C35 and C36 are usually glycine tRNA identity determinants (Giegé et al. 1998). It is not known, however, whether the SR1 GlyRS is also discriminative against C36. For mitochondrial tRNAs the identity determinants are largely unknown (Salinas-Giegé et al. 2015) and the origins of many of the tRNAs with altered anticodons have never been determined, so that these cases might also fit into the model as soon as more data become available. Nevertheless, the current data suggest a close connection between the tRNAs capturing a free codon and the respective aaRSs being able to correctly charge the cognate and the newly assigned tRNAs.

### Predictions based on the tRNA-loss driven codon reassignment theory

Our *tRNA loss driven codon reassignment* hypothesis presents several testable predictions that are mutually exclusive with the *codon capture* and *ambiguous intermediate* theories. We predict identification of (i) further yeast species with 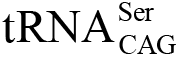 branching before the *SP* and *PC* split or within the group of *Pichia* and *Pachysolen* species, (ii) species with 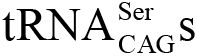 evolved from serine AGN-decoding tRNAs, and (iii) species with 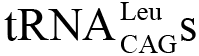 derived from tRNA_naaS_. Furthermore, we anticipate finding further yeasts with 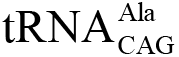 and species without 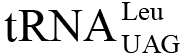 within the *PP* group. According to the *ambiguous intermediate* theory such findings would need to be treated as further independent ambiguous intermediate events. While this might be theoretically possible, it becomes increasingly unlikely that always the same codon is affected. Based on our assumption that tRNAs can only capture CUG codons if the respective aminoacyl-tRNA synthetases are non-discriminative against the mutated anticodon, we don't expect the CUG codon be captured by other tRNAs than leucine, serine and alanine encoding ones. The predictions of the *tRNA loss driven codon reassignment* model might best be tested by sequencing further yeast species.

## Materials and Methods

### Growth and lysis of *Pachysolen tannophilus NRRL Y-2460*

*Pachysolen tannophilus NRRL Y-2460* was obtained from ATCC (LGC Standards). Cells were grown in YFPD medium at 30°C, harvested by centrifugation (20' at 5,000 x g), and washed and resuspended in lysis buffer (50 mM HEPES pH 6,8, 100 mM KCl). The cells were disrupted by three passages through a French press (20,000 lb/in^2^) at 4°C, and intact cells and the cell debris removed by centrifugation (10' at 15,000 x g). The supernatant was subjected to SDS-PAGE gel electrophoresis.

### Genome annotation

The *Pachysolen* genome assembly (Liu et al. 2012) has been obtained from NCBI (GenBank accessions CAHV01000001-CAHV01000267). Gene prediction was done with AUGUSTUS (Stanke and Waack 2003) using the parameter “genemodel=complete”, the gene feature set of *Candida albicans*, and the standard codon translation table. The gene prediction resulted in 5,288 predicted proteins, out of which 4,210 contain at least one CUG codon. For mass spectrometry database search the database was multiplied so that each new database contains the CUG codons translated by another amino acid.

### Mass spectrometry analysis

SDS-PAGE-separated protein samples were processed as described by Shevchenko *et al.* (Shevchenko et al. 1996) The resuspended peptides in sample loading buffer (2% acetonitrile and 0.1% trifluoroacetic acid) were fractionated and analysed by an online UltiMate 3000 RSLCnano HPLC system (Thermo Fisher Scientific) coupled online to the Q Exactive HF mass spectrometer (Thermo Fisher Scientific). Firstly, the peptides were desalted on a reverse phase C18 precolumn (3 cm long, 100 μm inner diameter, 360μm outer diameter) for 3 minutes. After 3 minutes the precolumn was switched online with the analytical column (30 cm long, 75 μm inner diameter) prepared inhouse using ReproSil-Pur C18 AQ 1.9 μm reversed phase resin (Dr. Maisch GmbH). The peptides were separated with a linear gradient of 5-35% buffer (80% acetonitrile and 0.1% formic acid) at a flow rate of 300 nl/min (with back pressure 500 bars) over 90 min gradient time. The precolumn and the column temperature were set to 50°C during the chromatography. The MS data were acquired by scanning the precursors in mass range from 350 to 1600 m/z at a resolution of 70,000 at m/z 200. Top 30 precursor ions were chosen for MS2 by using data-dependent acquisition (DDA) mode at a resolution of 15,000 at m/z 200 with maximum IT of 50 ms. Data analysis and search were performed using MaxQuant v.1.5.2.8 as search engine with 1% FDR against the *Pachysolen* genome annotation database as annotated above, respectively. Search parameter for searching the precursor and fragment ion masses against the database were as described in Oellerich et al. (Oellerich et al. 2011) except that all peptides shorter than seven amino acids were excluded. To get confidence for the amino acids translated from CUG codons, we determined the observed fragment ions around each CUG-encoded residue. Only amino acids with fragment ions at both sides of the amino acid, which allow the determination of the mass of the respective amino acid, were regarded as supported by the data.

### tRNA phylogeny

tRNA genes in 60 sequenced yeast species and four *Schizosaccharomyces* species, which were used as outgroup (Table S1) (Mühlhausen and Kollmar 2014), were identified with tRNAscan (Lowe and Eddy 1997) using standard parameters. All genomes, in which tRNA_CAG_s were not found by tRNAscan, were searched with BLAST and respective tRNAs reconstructed manually. This especially accounts for the many tRNA_CAG_s having long introns (up to 287 bp). The intron-free tRNA_CAG_s were aligned against all *Candida albicans* cytoplasmic tRNAs to identify the closest related tRNA types for in-depth analysis. While the 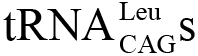 and 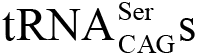 were easily identified, the *Pachysolen* tRNA_CAG_ grouped within the *Candida* alanine and valine tRNAs. To finally resolve tRNA codon type relationships and reconstruct tRNA_CAG_ evolution, we increased sequence and taxonomic sampling. Therefore, we randomly selected three to ten homologs from all leucine, serine, and alanine isoacceptor tRNAs from all 60 yeast species, and similar numbers of tRNAs from a selection of valine, phenylalanine, methionine, arginine, isoleucine and threonine codon types. Identical tRNA_CAG_ sequences from gene duplications were removed resulting in an alignment of 172 tRNA sequences (Fig. S2). To refine the resolution of tRNA relationships within codon family boxes, we manually removed mitochondrial Leu-, Ser-, and Ala-tRNAs from the datasets and performed separate phylogenetic analyses of all NAG-tRNAs (320 leucine isoacceptor tRNAs), NGA-tRNAs (776 serine isoacceptor tRNAs), and NGC-tRNAs (824 alanine isoacceptor tRNAs; Figs. S7-S9). Sequence redundancy was removed using the CD-HIT suite (Li and Godzik 2006) generating reduced alignments of representative sequences of less than 95% identity (80 Leu-tRNAs, 76 Ser-tRNAs, and 70 Ala-tRNAs).

Phylogenetic trees were inferred using Neighbour Joining, Bayesian and Maximum Likelihood based methods as implemented in ClustalW v.2.1 (Chenna et al. 2003), Phase v. 2.0 (Jow et al. 2002; Hudelot et al. 2003) and FastTree v. 2.1.7 (Price et al. 2010), respectively. The most appropriate model of nucleotide substitution was determined with JModelTest v. 2.1.5 (Darriba et al. 2012). Accordingly, FastTree was run with the GTR model for estimating the proportion of invariable sites and the GAMMA model to account for rate heterogeneity. Bootstrapping in ClustalW and FastTree was performed with 1,000 replicates. Phase was used with a mixed model, the REV-Γ model for the loops and the RNA7D-r model for the stem regions, which were given by a manually generated consensus tRNA secondary structure. Phase was run with 750,000 burnin and 3,000,000 sampling iterations, and a sampling period of 150 cycles. The phylogenetic network was generated with SplitsTree v.4.1.3.1 (Huson and Bryant 2006) using the Neighbor-Net method to identify alternative splits.

## Acknowledgments

We would like to thank Prof. Dr. Christian Griesinger for his continuous generous support, Dr. Hans-Dieter Schmitt for his help in growing *Pachysolen*, and Dr. Christof Lenz and Samir Karaca for help in analysing the mass spectrometry data. This project has been funded by a Synaptic Systems fellowship to SM.

## Author contributions

MK initiated the study. SM performed MS data and phylogenetic analyses. PF prepared experimental samples. UP performed MS experiments. HU was involved in MS data interpretation. MK assembled, aligned and analysed tRNA sequences. SM and MK wrote the manuscript.

